# Distinct infection and colonization strategies in cowpea leaves by three *Colletotrichum* species recently reported in Nigeria

**DOI:** 10.64898/2026.10.01.755927

**Authors:** Adenike O. Dada, Victor O. Dania, Olaniyi A. Oyatomi, Michael Abberton, Alejandro Ortega-Beltran

## Abstract

Cowpea, a crop of great importance in several regions in West Africa, faces many biotic and abiotic challenges. Several species of the genus *Colletotrichum* cause cowpea anthracnose disease (CAD), a major constraint to cowpea production severely affecting foliage and yield. Three *Colletotrichum* species—*C. cliviicola, C. truncatum*, and *C. chrysophilum* recently reported as causal agents of CAD in Nigeria were investigated in their infective processes and pathogenicity on cowpea leaves using histopathological and microscopic examination of stained leaf tissues. Inoculated leaves were examined at intervals to monitor fungal establishment, colonization, and reproductive development within host tissues. The three species completed the infection cycle, progressing from conidial germination and host penetration to tissue colonization, and sporulation. However, differences in both infective ability and pathogenicity were observed in representative isolates of each species. *C. truncatum* isolate was the most virulent, had extensive branching hyphae, produced severe tissue necrosis, abundant acervuli, and prominent seta formation. *C. cliviicola* also had substantial colonization and sporulation ability with dense conidial masses and mature acervuli. In contrast, *C. chrysophilum* had moderate infectivity with less extensive reproductive structures and tissue degradation. Here we provide the first histopathological report of infection and pathogenic development of these three species recently reported as causal agents of CAD.

## Introduction

Cowpea (*Vigna unguiculata* (L.) Walp) is a legume of great economic significance for several countries in sub-Saharan Africa (SSA), including Nigeria (Vanlauwe et al., 2019). However, the crop is constrained by several biotic and abiotic factors limiting its production and productivity. One of the most destructive fungal diseases affecting cowpea worldwide is anthracnose disease (CAD) (Pradhan et al., 2018; Satpathy and Beura, 2021). The disease, caused by several species of the genus *Colletotrichum*, can affect cowpea from seedling emergence to pod maturity, and rapidly develops under favorable environmental conditions (Pandey et al., 2023). CAD affects multiple aerial plant organs, including leaves, stems, petioles, flowers, and pods (Christopher et al., 2022). Given the critical role of cowpea as a source of food, nutrition, and income for millions of people, particularly in SSA, strategies to control CAD are required to achieving optimal productivity and food security (Gomes et al., 2019).

CAD disrupts normal physiological and metabolic processes, leading to significant reduction in content of proteins, carbohydrates, lipids, calcium, phosphorus, crude fibre, and ash in infected tissues, seeds, and husks (Amadioha and Enyiukwu, 2019). CAD outbreak can result in yield losses of >50% when effective management practices are not implemented (Dabbas et al., 2015). In addition, CAD reduces marketability and economic value of cowpea through quality deterioration, impaired seed development, and contamination with harmful fungal metabolites (Gloria and Ataga, 2021; Viljoen et al., 2025).

The broad geographical occurrence of CAD is largely associated with the seed-borne nature of several *Colletotrichum* species, which facilitates long-distance dissemination of inoculum through infected planting materials (Awurum and Ucheagwu, 2013; Pandey et al., 2023). Infected seeds serve as important survival structures, enabling CAD pathogens to initiate new infections after planting provided that conducive environmental conditions exist (Yadav et al., 2025). In addition to seed transmission, *Colletotrichum* species can survive in infected crop residues and plant debris, where they remain viable for extended periods and can act as primary sources of inoculum in subsequent cropping seasons (Geetha et al., 2013; Usman et al., 2025). Following primary infection, abundant conidia are produced in acervuli on diseased tissues and disseminated mainly by rain splash, irrigation water, and wind-driven rain, resulting in rapid secondary spread of the disease within and among fields (De Silva et al., 2017; Usman et al., 2025). Successful infection begins with conidial adhesion and germination on the leaf surface, followed by appressorium-mediated penetration and subsequent colonization of host tissues through a hemibiotrophic infection strategy characteristic of many *Colletotrichum* species (Boufleur et al., 2021). The polycyclic nature of CAD, coupled with favorable conditions, contributes significantly to disease establishment in susceptible cowpea germplasm.

In Nigeria, *C. lindemuthianum* and *C. destructivum* were historically reported as the principal causal agents of CAD based largely on morphological characterization (Latunde-Dada, 1997; Dania and Gbadamosi, 2019; Enyiukwu et al., 2020). However, recent molecular investigations have revealed other *Colletotrichum* spp. associated with CAD. The fungi, *C. cliviicola, C. siamense, C. chrysophilum*, and *C. truncatum* were reported for the first time, causing CAD in Nigeria recently (Adeyeye et al., 2024; Dada et al. 2023, 2025, 2026). While previous studies have characterized the diversity, pathogenicity, and host responses of *Colletotrichum* species associated with CAD in Nigeria, the infection processes and tissue colonization patterns of the recently identified species remain poorly understood. In particular, it is not known whether these *Colletotrichum* species differ in their infection strategies, rate of tissue colonization, or progression within cowpea tissues. Such differences may provide important insights into pathogen–host interactions and help explain variations in disease development and pathogenicity among *Colletotrichum* species associated with CAD. Therefore, the current study aimed to characterize and compare the infection processes and tissue colonization patterns of three *Colletotrichum* species associated with CAD in Nigeria, providing further insight into their infection strategies and interactions with cowpea tissues. The results will allow designing targeted management strategies, including agronomic practices, bioprotectant usage, and host resistance breeding.

## Materials and methods

### Sowing of seeds and leaf preparation

In a screenhouse, three 10-kg pots containing sterilized soil were sown with two seeds of the CAD susceptible cowpea variety, Ife Brown. Seedlings were thinned to one plant per pot at seven days after sowing. The plants were maintained under screenhouse conditions for three weeks with adequate watering before leaves were harvested for detached leaf assays (DLA) as previously described (Dada et al. 2025). Collected leaves were surface-sterilized in 1% sodium hypochlorite (NaOCl) and rinsed in three changes of sterile distilled water and dried with sterile paper towel in a biosafety cabinet. Plastic crisper boxes (31 × 23 × 10 cm) were sterilized with 2% NaOCl, rinsed with two changes of sterile distilled water, and drained under the biosafety cabinet. Cotton wool and paper towels wrapped in aluminum foil were autoclaved (121°C, 15 min) and laid in equal proportion in the boxes. Fifteen ml sterile water was evenly spread in each laid box to keep it moist and three surface-sterilized leaves were placed in the box with the abaxial sides facing the lid.

### Source of isolates and pathogen characterization

Previously characterized isolates of *C. chrysophilum* (isolate CC1_NG) and *C. cliviicola* (isolate CC24_NG) (Dada et al., 2023; 2025) were obtained from the fungal collection of the Pathology and Mycotoxin Laboratory of IITA. Isolates of *C. truncatum* were obtained from naturally infected cowpea plants collected from the IITA-Ibadan research fields. All isolates had the distinct falcate-shape spore belonging to the *C. truncatum* group. Representative isolate (CC18_NG) was further characterized based on molecular analysis. PCR amplification was performed using the primer pairs ACT512 (ATGTGCAAGGCCGGTTTCGC) and ACT783 (TACGAGTCCTTCTGGCCCAT), targeting the actin (ACT) gene, and GDF (GCCGTCAACGACCCCTTCATTGA) and GDR (GGGTGGAGTCGTACTTGAGCATGT), targeting the glyceraldehyde-3-phosphate dehydrogenase (GAPDH) gene. The resulting amplicon was purified and subjected to Sanger sequencing using an ABI 3130xl Genetic Analyzer (Applied Biosystems). The nucleotide sequences obtained from the query isolate CC18_NG were assembled using BioEdit and verified via the BLAST searches against the NCBI database. The sequences were deposited in the NCBI database under accession numbers OR965855 (actin) and OR965854 (GADPH). The actin and GAPDH sequences of CC18_NG were aligned alongside 19 reference sequences of *Colletotrichum* spp. including fungi previously characterized by Dada et al., (2025) as in Table 1. The sequences were trimmed to uniform lengths, concatenated, and used to infer a single phylogenetic tree using the Maximum Likelihood method with the Kimura-2-parameter model in Molecular Evolutionary Genetics Analysis (MEGA) v12.

**Table 1.**
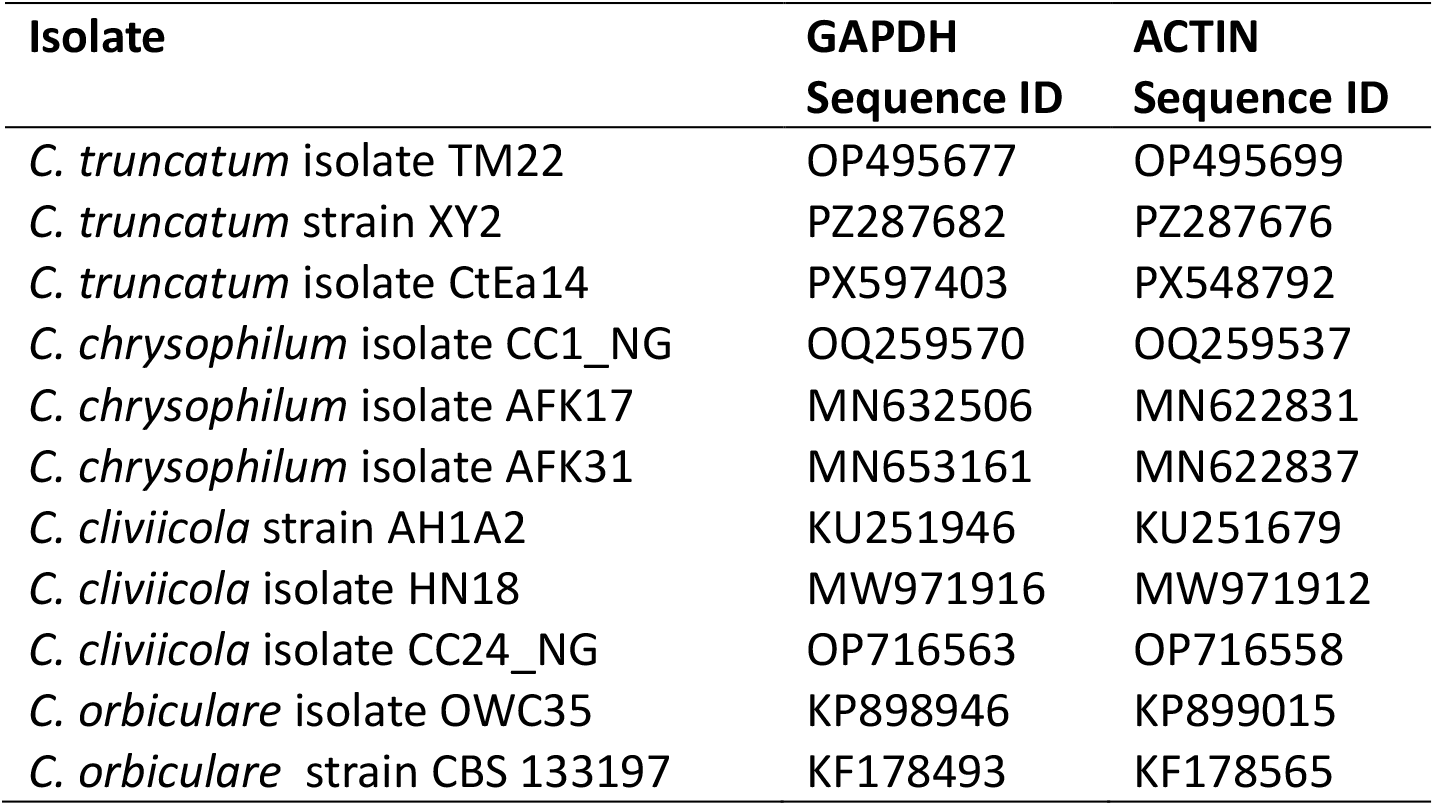
Details of isolates used for phylogenetic analysis.

| Isolate | GAPDH<br>Sequence ID | ACTIN<br>Sequence ID |
| --- | --- | --- |
| *C. truncatum* isolate TM22 | OP495677 | OP495699 |
| *C. truncatum* strain XY2 | PZ287682 | PZ287676 |
| *C. truncatum* isolate CtEa14 | PX597403 | PX548792 |
| *C. chrysophilum* isolate CC1_NG | OQ259570 | OQ259537 |
| *C. chrysophilum* isolate AFK17 | MN632506 | MN622831 |
| *C. chrysophilum* isolate AFK31 | MN653161 | MN622837 |
| *C. cliviicola* strain AH1A2 | KU251946 | KU251679 |
| *C. cliviicola* isolate HN18 | MW971916 | MW971912 |
| *C. cliviicola* isolate CC24_NG | OP716563 | OP716558 |
| *C. orbiculare* isolate OWC35 | KP898946 | KP899015 |
| *C. orbiculare* strain CBS 133197 | KF178493 | KF178565 |

### Inoculum preparation and pathogen inoculation

Inoculum suspensions were prepared from two representative isolates each of *C. chrysophilum, C. cliviicola*, and *C. truncatum* grown in potato dextrose agar medium for 2 days at 28°C. Spore concentrations were adjusted to 10^6^ spores/mL. TWEEN^®^20 was added at 1 mL/L as a surfactant to facilitate uniform distribution of conidia on the surface of leaves to be inoculated. The leaves were arranged on the moist paper towels in the crisper boxes, and a tiny hole was created in three angles on each leaf with a sterile needle. Then, an aliquot of 40 µL inoculum was applied to the spots using a micropipette, following the procedure of Dada et al. (2025). Each isolate constituted a treatment, with three detached leaves serving as replicates such that each crisper box had a single isolate treatment in three angles of each of three leaves (Fig. 1). Treatments were arranged in a completely randomized design. Control leaves were treated with sterile water. The crisper boxes were subsequently covered and sealed with cling film to prevent contamination and maintain high humidity conditions.

**Figure 1.**
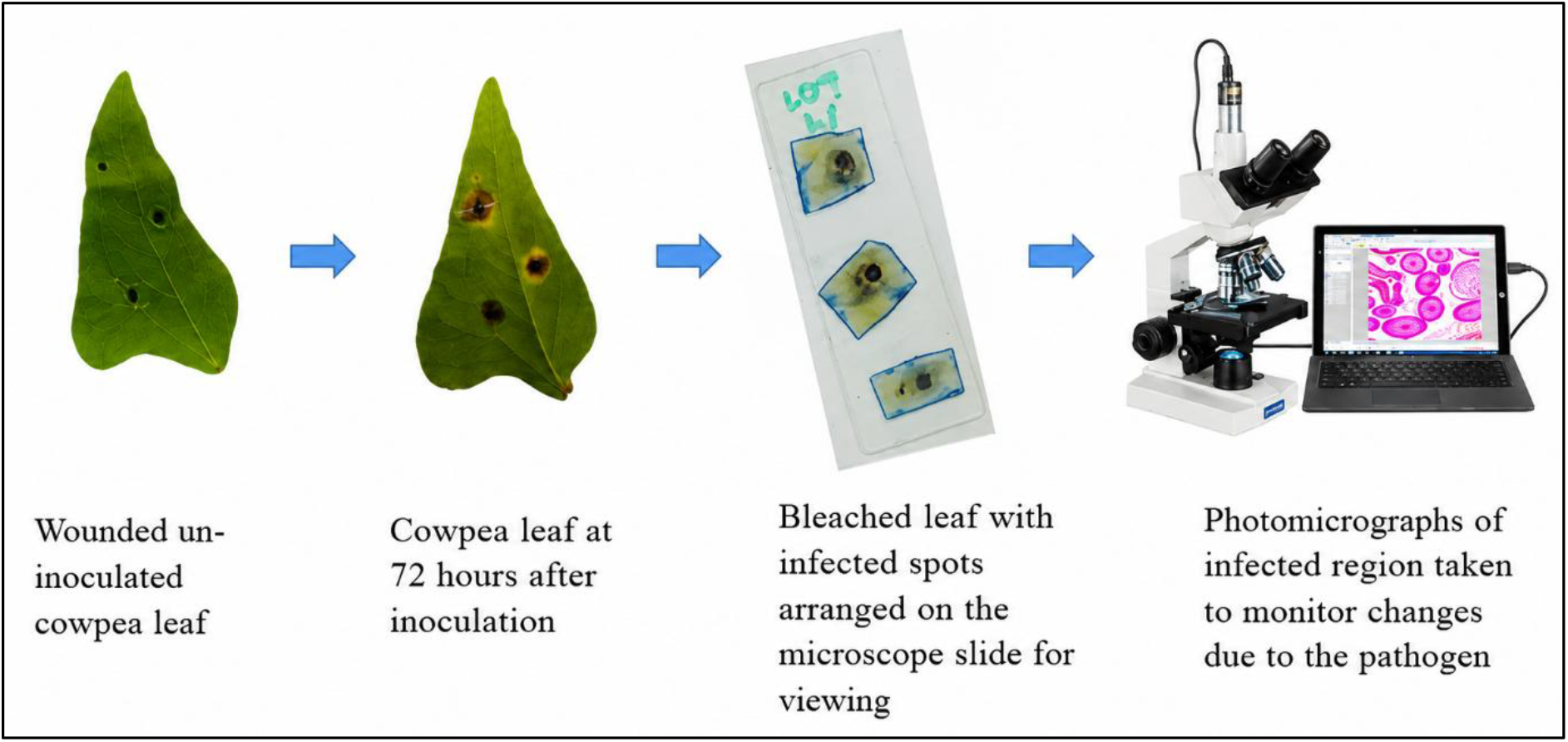
Procedure for infective studies of *Colletotrichum* species in Ife-Brown cowpea variety.

### Staining and microscopy

Staining and microscopy procedures were carried out according to the method of Latunde-Dada et al. (2007), as illustrated in Fig. 1. Sections were excised from the inoculated portions of the leaves at 24, 48, and 72 hr after inoculation (HAI) and decolorized for 12 hr in 0.15% (w/v) trichloroacetic acid prepared in a 3:1 (v/v) ethanol–chloroform mixture. The decolorized tissues were subsequently stained for 2 hr in 0.025% (w/v) aniline blue solution, placed on a microscope slide and mounted on a compound microscope for examination.

### Data collection and statistical analyses

The lesion sizes were measured and photomicrographs of infected tissues were taken at different observation periods: 24, 48, and 72 HAI. Infection structures and tissue colonization patterns included on conidial germination, appressorium formation, penetration events, intracellular colonization and sporulation at three observation periods. Numeric data were analyzed using R software v4.5.3.

## Results

### Characterization of *C. truncatum* isolate

BLAST search conducted with actin and GAPDH sequences of the isolate revealed >95% identity to *C. truncatum* isolates deposited in NCBI. Phylogenetic analysis (Fig. 2) using these two loci confirmed high level of similarity with reference *C. truncatum* isolates at >96% bootstrap value. All *C. truncatum* isolates were clearly distinguished from other species.

**Figure 2.**
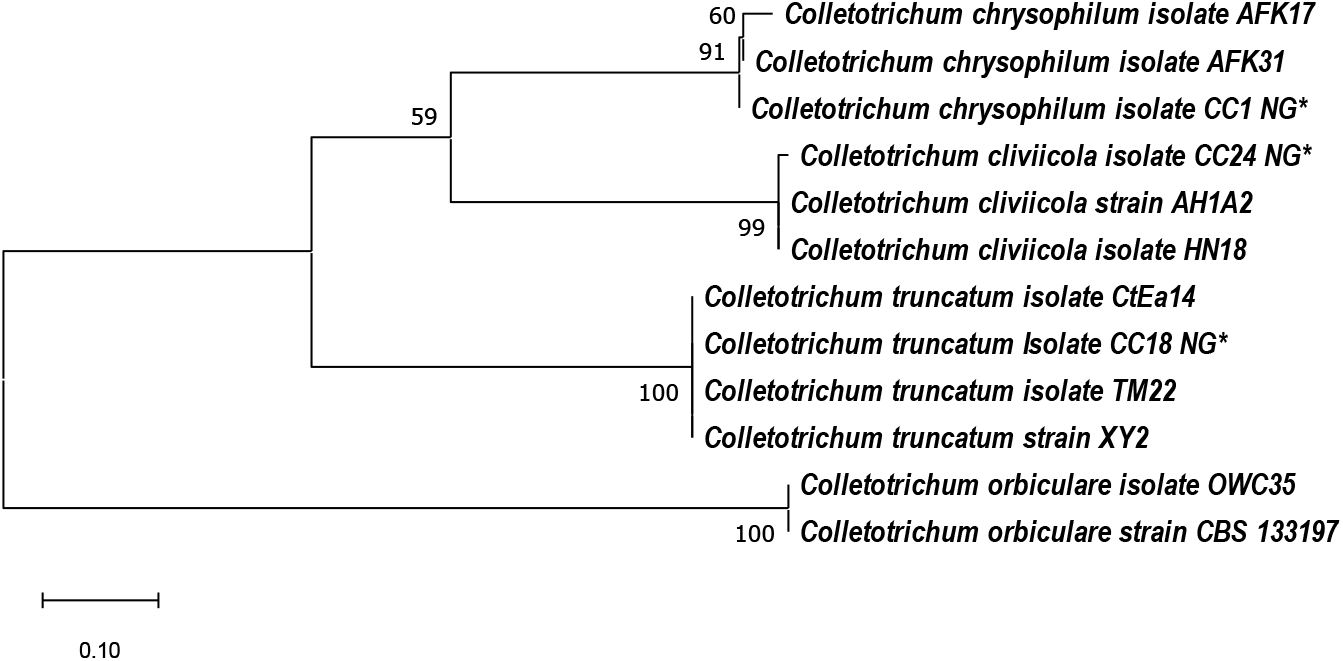
Phylogenetic relationship of *Colletotrichum* species causing cowpea anthracnose disease in cowpea leaves. Names of the isolates used in the current study include an asterisk (*). *Colletotrichum orbiculare* species are included as an outgroup

### Symptom development on cowpea leaves

All inoculated leaves developed characteristic CAD symptoms, whereas leaves treated with sterile distilled water remained symptomless throughout the experiment (Fig. 3). Initial symptoms appeared as small reddish-brown necrotic lesions surrounded by distinct chlorotic halos. As infection progressed, lesions enlarged and coalesced, resulting in necrosis.

**Figure 3.**
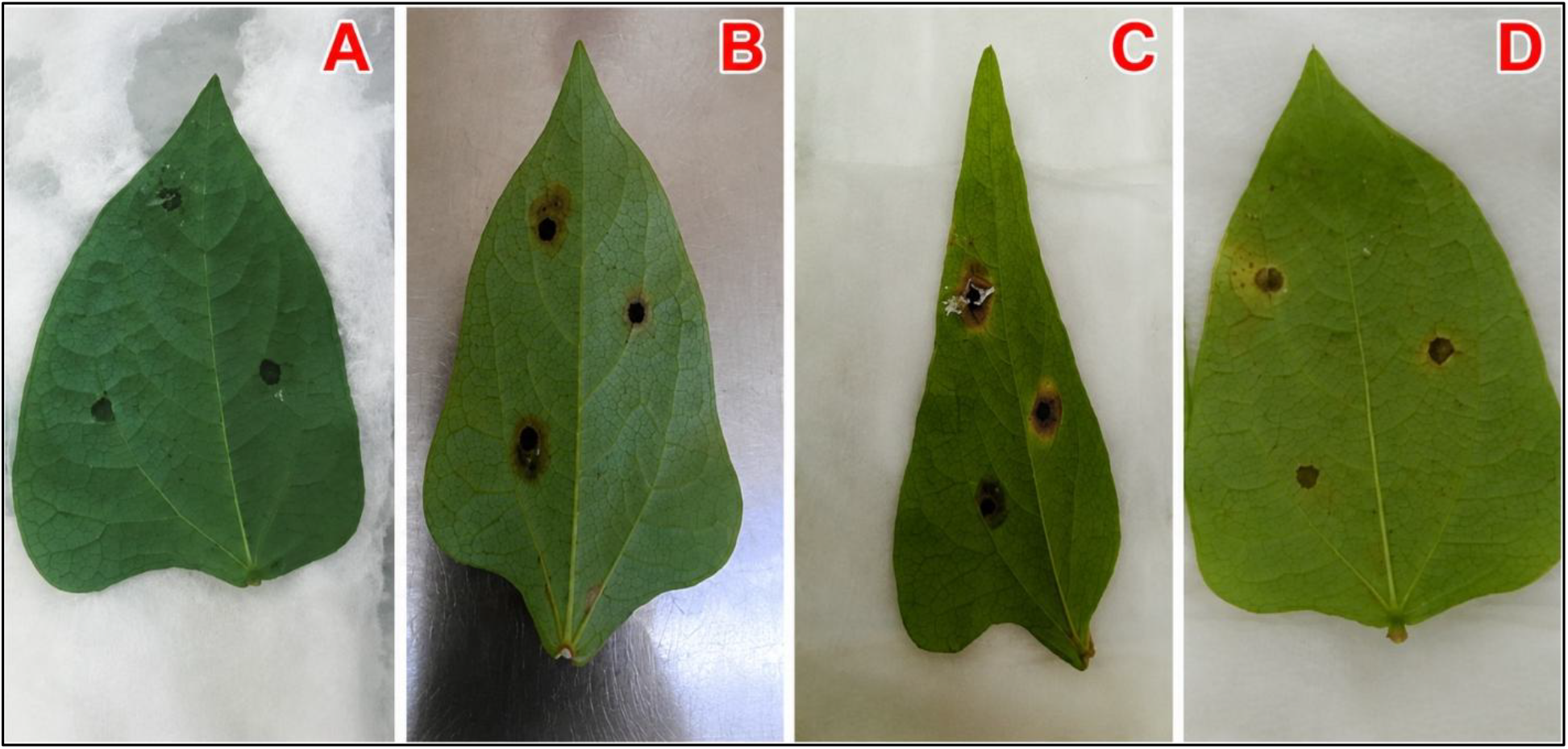
Ife Brown cowpea leaves inoculated with sterile distilled water (A), *Colletotrichum chrysophilum* (B), *C. truncatum* (C), and *C. cliviicola* (D) at 72 hours after inoculation.

The lesion size differed among the three *Colletotrichum* species with time (P<0.001) (Table 2). At 72 HAI, leaves inoculated with *C. truncatum* developed larger lesions of about 4.40±0.75 mm (Table 3) with more pronounced chlorosis and extensive tissue necrosis than those inoculated with *C. chrysophilum* (4.20±0.65 mm) and *C. cliviicola* (2.77+0.40 mm) (Fig. 4).

**Figure 4.**
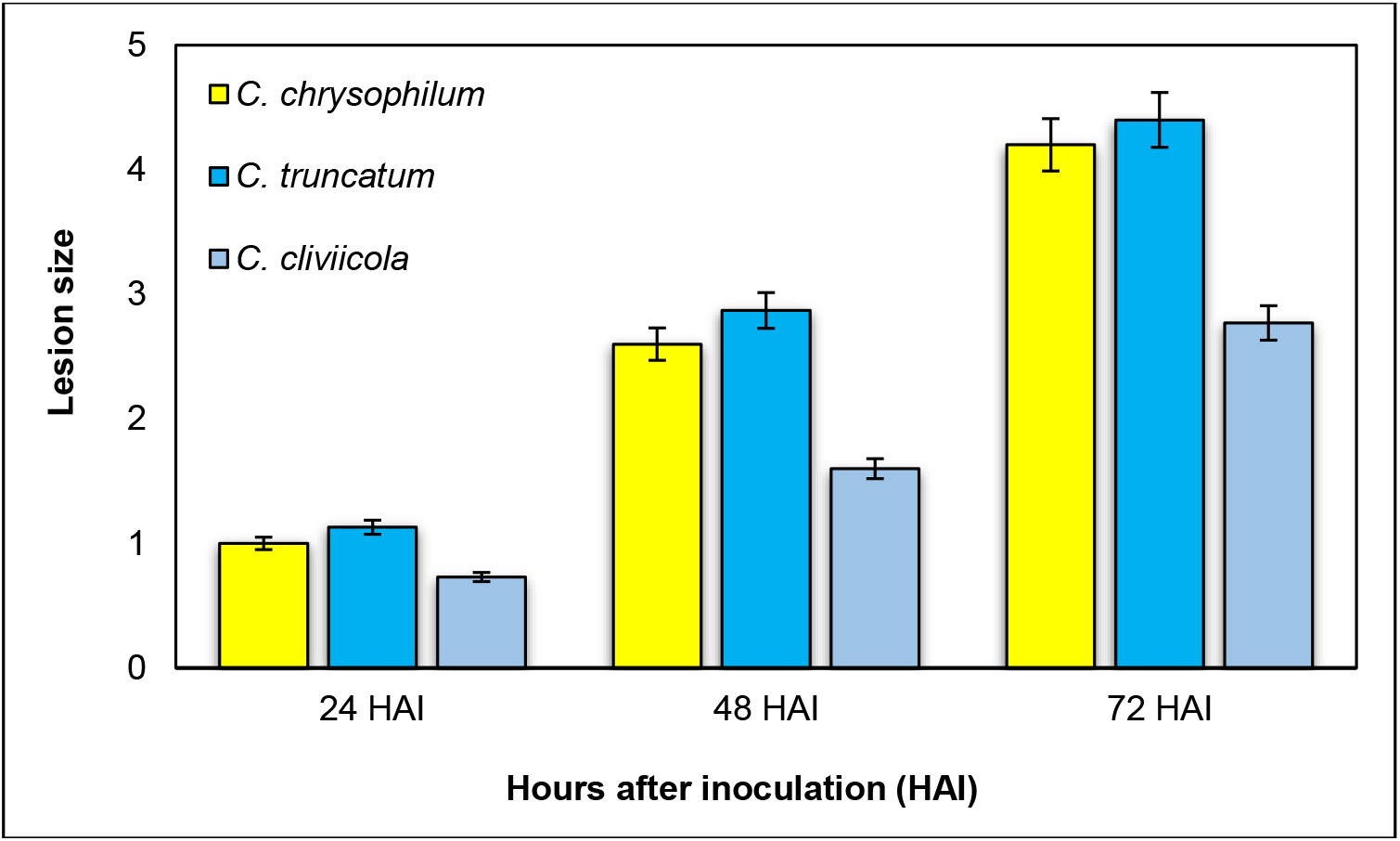
Comparison of lesion size produced on produced on cowpea leaf by *Colletotrichum chrysophilum, C. cliviicola* and *C. truncatum* at 24, 48 and 72 hours after inoculation (HAI)

### Histopathological development of *Colletotrichum* species during infection of cowpea leaves

Microscopic examination demonstrated that the three *Colletotrichum* species had a common pattern of infection series on cowpea leaves, progressing through conidial germination, appressorium formation, host penetration, hyphal colonization, acervulus development, and sporulation (Figs. 5-7). Nevertheless, clear differences were observed in the rate of development, extent of host colonization and reproductive capacity among the three species.

**Table 2.** Mean square of anthracnose lesion produced on cowpea leaf by *Colletotrichum chrysophilum, C. cliviicola*, and *C. truncatum* at 24, 48 and 72 hours after inoculation (HAI)

| Source of variation | df | Mean Square | F-value | P-value |
| --- | --- | --- | --- | --- |
| Lesion sizes | 2 | 3.090 | 15.51 | <0.001 |
| Time (HAI) | 2 | 18.064 | 90.65 | <0.001 |
| Lesion size × HAI | 4 | 0.377 | 1.89 | 0.156 |
| Error | 18 | 0.199 |  |  |

**Figure 5.**
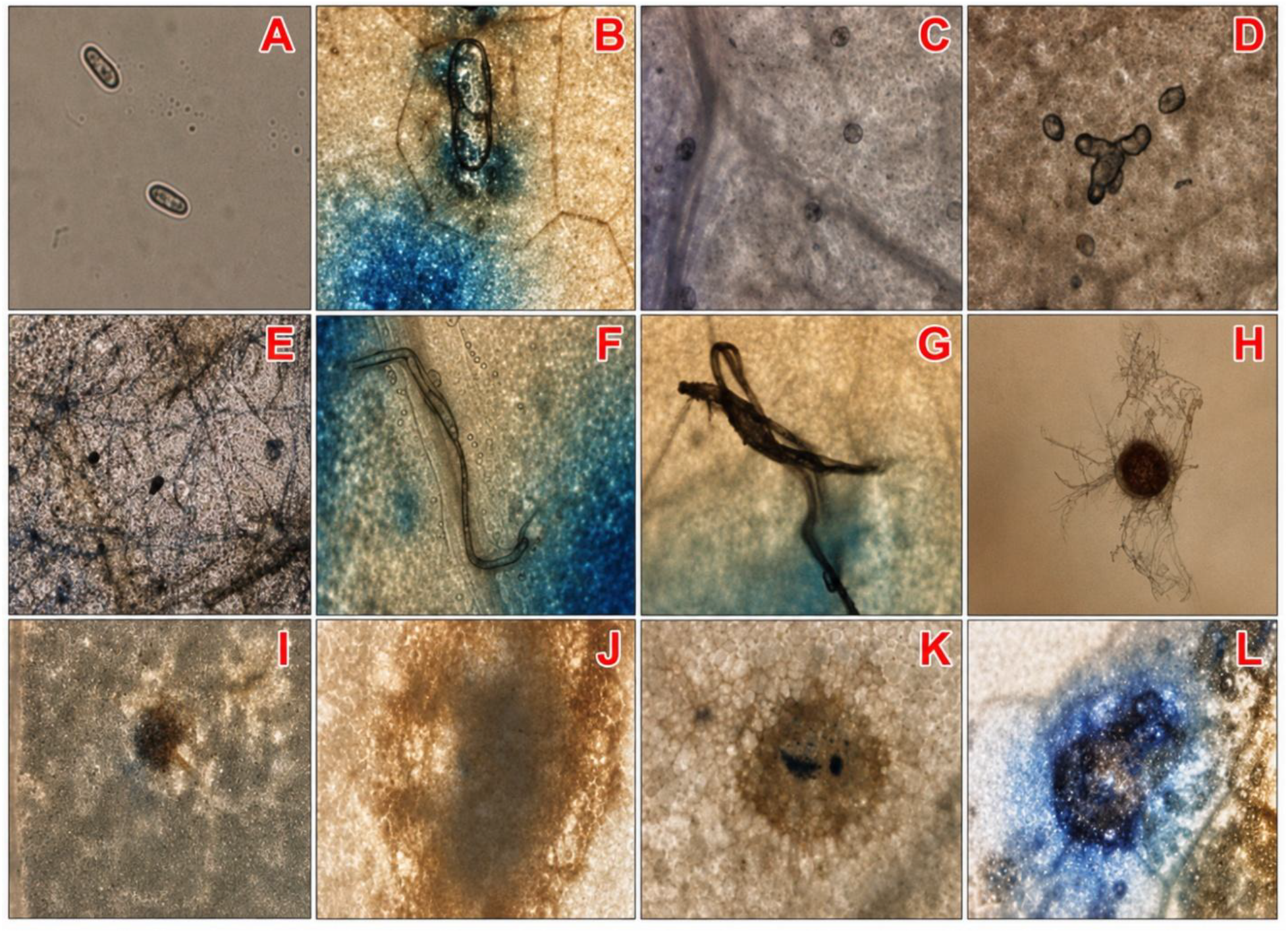
Microscopic observations of bleached cowpea leaf tissues inoculated with *Colletotrichum chrysophilum* showing different stages of fungal development and host colonization. Conidia(A); germinating spores and early penetration structures (B-D); hyphal colonization(E-G); developing acervuli and host tissue responses (H-K); mature sporulating structure/acervulus (L).

**Figure 6.**
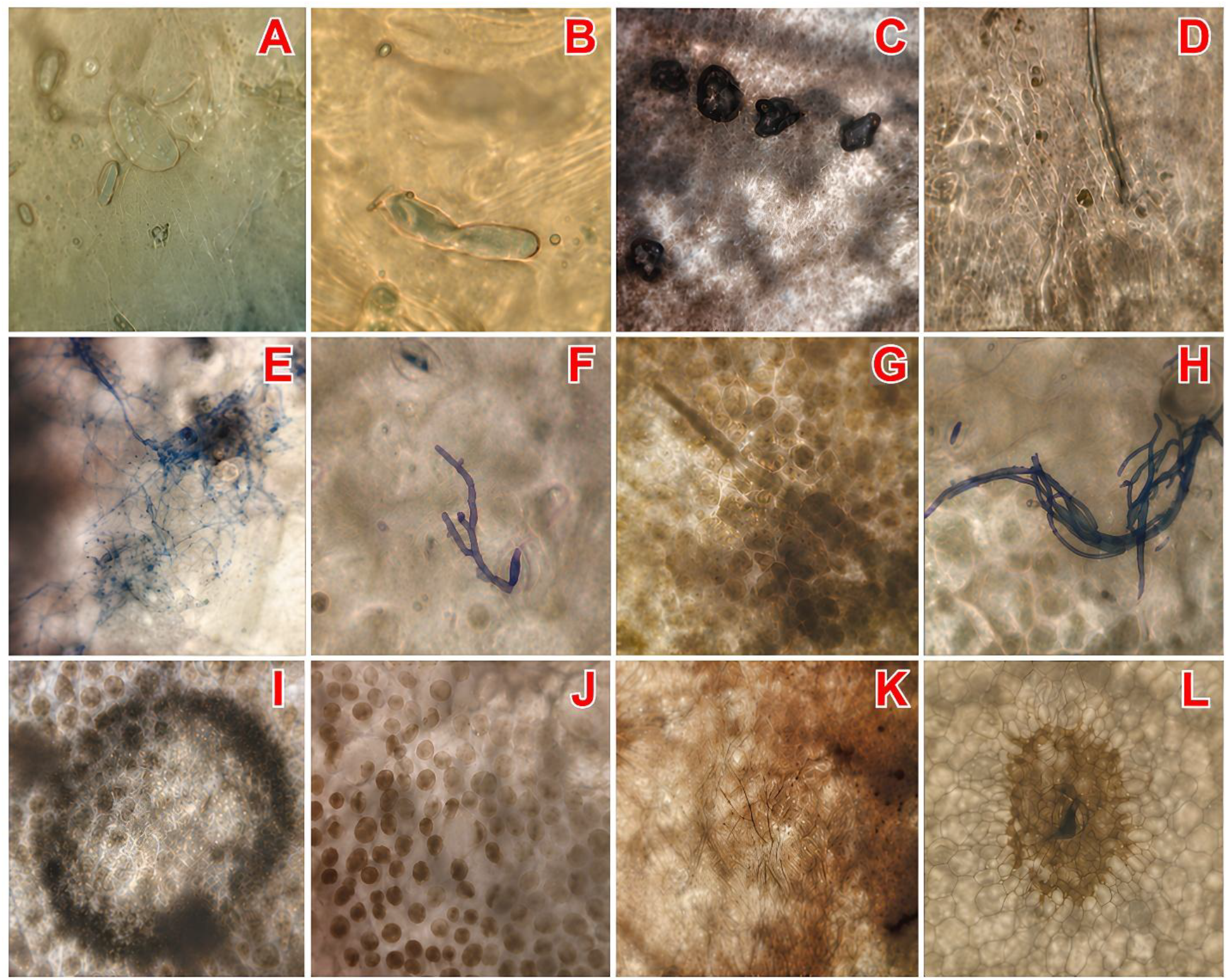
Microscopic observations of bleached cowpea leaf tissues infected with *Colletotrichum cliviicola* showing fungal developmental stages and reproductive structures. Conidia (A); early penetration and germination structures (B–D); branching and invasive hyphae (E–F); aggregated fungal mass and setae (G); acervuli/conidial bodies (H, L); tissue colonization and necrosis (I–J); setae associated with acervuli (K).

**Figure 7.**
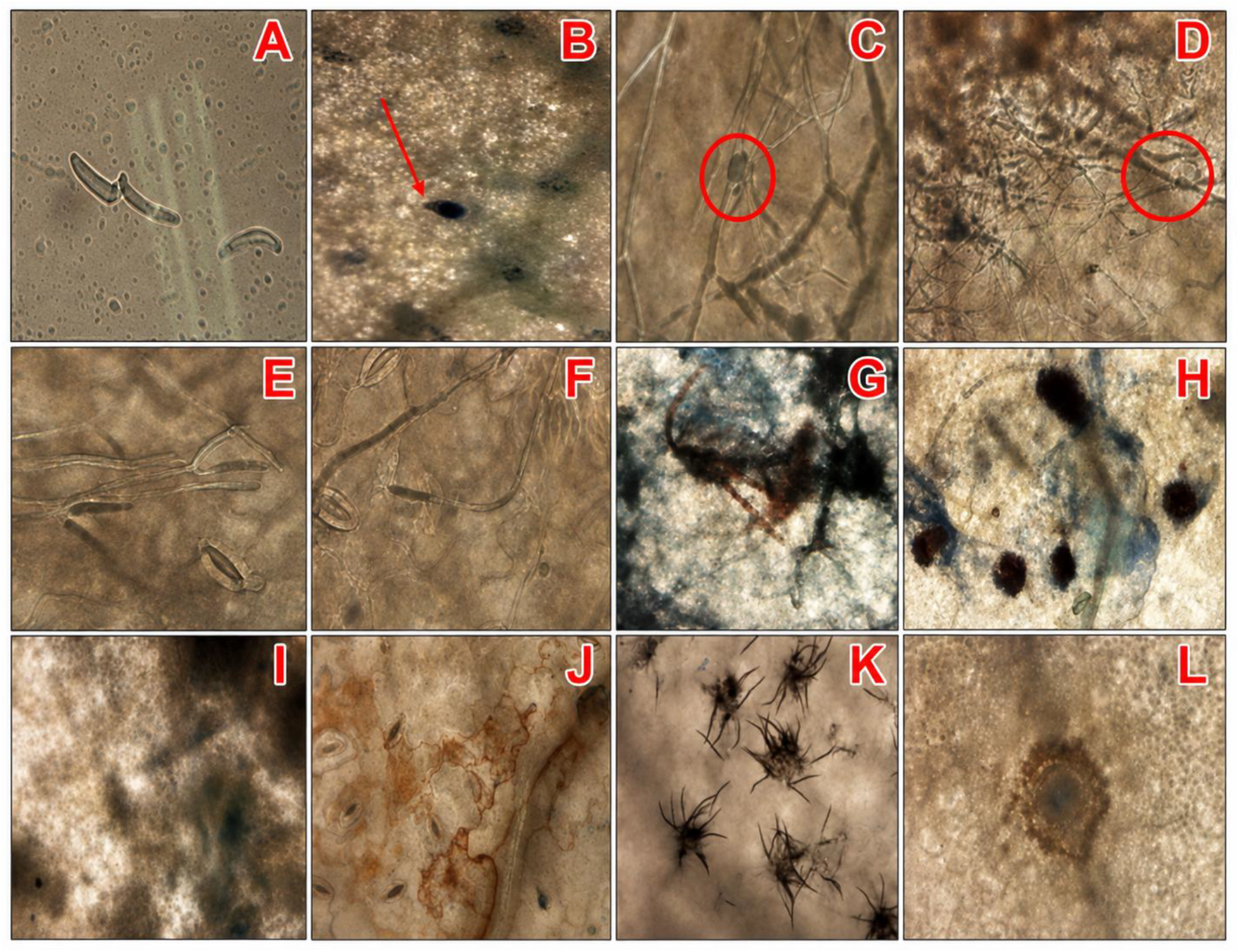
Microscopic examination of bleached cowpea leaf tissues infected with *Colletotrichum truncatum* showing fungal developmental stages and reproductive structures. Conidium(A); germinating conidium (B); melanized infection structures (C); invasive and branching hyphae (D–F); colonized host tissue (G); appressorium-like structure (H); mature acervuli (I, L); conidial masses (J); necrotic lesion center (K).

### Conidial germination and appressorium formation

The infection process began with the deposition of conidia on the leaf surface. Conidia of *C. chrysophilum* and *C. cliviicola* were hyaline, aseptate and ovoid, whereas those of *C. truncatum* were characteristically hyaline, falcate, and uniseptate (Figs. 5A, 6A, 7A). Germination commenced within 24 HAI, producing germ tubes that elongated over the epidermal surface before differentiating into irregularly shaped, dark colour appressoria (Figs. 5B-D, 6B-D, 7B-C). These melanized infection structures marked the beginning of host penetration and indicated successful establishment of the pathogen.

### Penetration and hyphal colonization

Following appressorium formation, penetration hyphae developed and entered the epidermal tissues. Secondary hyphae subsequently emerged and spread through adjacent host cells, producing extensive intracellular and intercellular colonization (Figs. 5E-G, 6E-G, 7D-G). The secondary hyphae were thinner than the primary hyphae and possessed narrow pointed tips that facilitated further invasion of surrounding tissues. Progressive branching resulted in dense mycelial networks that occupied increasing proportions of the infected tissues. As colonization advanced, host cells gradually lost their structural integrity, and extensive tissue collapse became evident.

Although the sequence of infection was similar among the three species, there were differences in colonization intensity. *C. chrysophilum* exhibited comparatively moderate hyphal development with less extensive invasion of host tissues (Fig. 5E-G). In contrast, *C. cliviicola* produced dense branching hyphae and conspicuous fungal aggregates that occupied larger areas of the infected tissue (Fig. 6E-G). The most extensive colonization was observed in *C. truncatum*, where rapidly expanding hyphal networks were associated with severe disruption of host tissues and early development of necrotic lesions (Fig. 7D-G).

### Host tissue responses

Host responses became increasingly evident as fungal colonization progressed. Infected cells exhibited progressive discolouration, degeneration and collapse, culminating in the formation of reddish-brown necrotic lesions. These symptoms were caused more rapidly by *C. truncatum*, which were clearly visible by 48 HAI. In contrast, leaves inoculated with *C. chrysophilum* and *C. cliviicola* required a longer period before comparable necrotic symptoms became fully expressed. Tissue degradation was consistently more extensive in leaves inoculated with *C. truncatum*, followed by *C. cliviicola*, whereas *C. chrysophilum* caused comparatively less damage.

### Acervulus development and sporulation

As colonization progressed, compact fungal aggregates differentiated into immature acervuli beneath the epidermis (Figs. 5H-K, 6G-L, and 7H-L). These reproductive structures enlarged with time and eventually ruptured the host epidermis to produce mature acervuli containing abundant conidia. Orange to golden-brown conidial masses exuded from mature acervuli, indicating completion of the infection cycle and initiation of secondary inoculum production.

Dark brown septate setae were observed in association with mature acervuli of both *C. truncatum* and *C. cliviicola*, although these structures were more numerous, longer and more conspicuous in *C. truncatum. C. chrysophilum* also produced reproductive structures but with comparatively fewer and less developed acervuli. *C. truncatum* exhibited the highest reproductive capacity, characterized by abundant acervuli, profuse conidial masses and extensive seta formation. *C. cliviicola* displayed a similar developmental pattern but with slightly reduced reproductive intensity, whereas *C. chrysophilum* showed comparatively moderate sporulation and tissue colonization than *C. truncatum*.

## Discussion

The ability of *Colletotrichum* species to produce reproductive structures on infected host tissues has important implications for the epidemiology and persistence of cowpea anthracnose disease. Once infection is established, diseased plant tissues may not only reflect the consequences of pathogen colonization but may also become active sources of inoculum that sustain subsequent cycles of infection. The infection process observed in the current study agrees with the general pathogenic behavior that has been previously reported for *Colletotrichum* spp., which typically involves attachment of conidia to the host surface, germination, appressorium-mediated penetration, intracellular or subcuticular colonization, and eventual sporulation on necrotic tissues. The presence of germinating conidia and elongated germ tubes in the infected tissues suggests that the isolates were able to recognize and adapt to the cowpea leaf surface environment, thereby initiating successful infection. The formation of invasive hyphae and branching mycelial networks further indicates active nutrient acquisition and spread within host tissues, which are characteristic features of anthracnose development in various crops (Bailey et al., 1990; Latunde-Dada et al., 1997; Smith et al., 1999; O’Connell et al., 2000; da Silva et al., 2020; Usman et al., 2026). Among the three examined species, *C. truncatum* exhibited comparatively more aggressive infective behavior, as evidenced by extensive mycelial proliferation, pronounced tissue necrosis, abundant acervuli formation, and conspicuous setae development. *C. truncatum* has been reported to cause anthracnose in several economically important pulse crops, including sword bean (Shi et al., 2022), lima bean (Brito et al., 2025), mung bean (Singla et al., 2024) and soybean (Mandloi et al., 2025). More importantly, yield losses of up to 100% by *C. truncatum* have been reported in unmanaged soybean and lima bean crops (Gomes and Nascimento, 2018; Dias et al., 2019). Comparatively, the high density of reproductive structures observed in tissues infected by *C. truncatum* in the present study may be attributed to greater pathogenic fitness and reproductive efficiency on cowpea leaves. This observation is consistent with previous reports identifying *C. truncatum* as a major causal agent of CAD in Nigeria and an economically important pathogen capable of causing severe yield losses under favorable environmental conditions (Adeyeye et al., 2024). The prominent seta formation associated with *C. truncatum* further supports its identity as a typical anthracnose pathogen and reflects advanced reproductive development within infected tissues (Shahriar et al., 2023; Zhang et al., 2024).

Although *C. chrysophilum* and *C. cliviicola* also demonstrated successful infection and colonization of cowpea leaves, as previously confirmed by Dada et al. (2023, 2025), differences were observed in the extent of tissue destruction and sporulation intensity. *C. chrysophilum* exhibited relatively moderate pathogenicity, characterized by successful conidial germination and hyphal establishment but comparatively fewer reproductive structures and less severe tissue degradation. The pathogen appeared to have a slower colonization rate or lower adaptation efficiency on cowpea tissues compared with *C. truncatum*. This finding is similar to the report of Liu et al. (2019), stating that *C. truncatum* was the most aggressive *Colletotrichum* species affecting cassava in China whereas *C. chrysophilum* showed more virulence than *C. truncatum* isolate in some cassava cultivars suggesting that the crop response to attack by each of the pathogen species may be due to variability in their susceptibility levels (Damascena et al., 2026).

*C. cliviicola* showed substantial infective ability, with abundant hyphal aggregation, mature acervuli, and dense conidial masses, indicating effective colonization and sporulation capacity. However, seta formation was less conspicuous in the pathogen than in *C. truncatum*, suggesting possible species-specific differences in reproductive morphology and infection strategy. Nevertheless, the study confirmed its ability to infect cowpea by showing evidence of structures that have been previously identified in the *C. cliviicola’s* infectivity process (Zhang et al., 2023). The variation in infective behavior among the three *Colletotrichum* spp. may be attributed to differences in virulence factors and reproductive efficiency. Anthracnose pathogens are known to produce cell wall-degrading enzymes, toxins, and melanized infection structures that facilitate penetration and tissue colonization (Remli et al., 2017; Ntui et al., 2021; Jeyaraj et al., 2023). Therefore, *C. truncatum* may have greater capacity for enzymatic degradation in cowpea than the two other evaluated species. Additionally, differences in host-pathogen compatibility may have influenced the level of colonization achieved by each species.

## Conclusion

The ability of *Colletotrichum cliviicola, C. chrysophilum*, and *C. truncatum* to infect cowpea through similar infection stages but with distinct patterns of host colonization, tissue degradation, and reproductive development highlights the importance of considering multiple *Colletotrichum* species in the epidemiology and management of cowpea anthracnose. The greater virulence and extensive tissue colonization, acervuli formation, and setae production observed in *C. truncatum*, alongside the substantial colonization and sporulation capacity of *C. cliviicola*, suggest that these species may differ in their contribution to disease development and epidemiology. These findings demonstrate that pathogenicity vary among *Colletotrichum* species associated with CAD highlighting the importance of considering species level variation when evaluating disease development, host responses and when developing strategies for anthracnose resistance and disease management in cowpea.

## Data Availability Statement

The datasets used and/or analyzed during the current study are available from the corresponding authors on reasonable request.

## Acknowledgments

The authors are grateful to all members of the Genetic Resource Centre and Pathology and Mycotoxin Unit of IITA that contributed to the execution of this study.

## Author Contributions

A.O.D., V.O.D, and A.O.-B. designed the experiments. A.O.D. performed the experiments. A.O.-B., O.A.O. and M.A. provided materials and supervision. A.O.D. wrote the manuscript. V.O.D., O.A.O., M.A., and A.O.-B. edited the manuscript. All authors gave final approval to publish the manuscript

